# Modifications of lipoic arm by reactive nitrogen species regulate α-ketoacid dehydrogenases

**DOI:** 10.1101/2022.01.31.478543

**Authors:** Gretchen L. Seim, Zixiang Fang, David J. Pagliarini, Jing Fan

## Abstract

Mitochondrial α-ketoacid dehydrogenases, including the pyruvate dehydrogenase complex (PDHC) and the oxoglutarate dehydrogenase complex (OGDC), are a family of multi-subunit enzyme complexes that use a lipoic arm to transfer an acyl group to coenzyme A (CoA). The regulation of α-ketoacid dehydrogenases plays crucial roles in mitochondrial metabolism and cellular energy homeostasis. We previously found that PDHC and OGDC become profoundly inhibited in macrophages upon classical activation, causing substantial remodeling of the TCA cycle. This inhibition was driven by the loss of the catalytically active lipoic moiety; however, the molecular mechanism causing this loss was not clear. Here we show that reactive nitrogen species (RNS), which are produced by activated macrophages, can cause a series of thiol-modifications to the lipoic arm that inactivate PDHC and OGDC. CoA-SNO, the non-enzymatic product between RNS and the E2 subunit’s natural substrate CoA, plays a key role in efficiently delivering RNS mediated modifications onto the lipoic arm. This work reveals a new biochemical mechanism capable of substantially regulating mitochondrial α-ketoacid dehydrogenases, which has potential relevance for a range of physiological and pathological conditions.

## Introduction

Mitochondrial α-ketoacid dehydrogenases are a family of multi-subunit enzyme complexes that includes pyruvate dehydrogenase complex (PDHC), oxoglutarate dehydrogenase complex (OGDC), and branched-chain α-ketoacid dehydrogenase complex (BCKDC). These enzymes function at critical crossroads in the metabolic network, controlling carbohydrate and amino acid metabolism, mitochondrial energy production, and cellular redox state.^1^ Specifically, PDHC catalyzes the conversion of pyruvate to acetyl-CoA, and OGDC catalyzes the oxidation of α-ketoglutarate to succinyl-CoA; thus, both control the entrance of major carbon sources into the TCA cycle. α-Ketoacid dehydrogenase complexes use similar catalytic mechanisms involving coupled reactions with three subunits.^1–4^ The E1 subunit decarboxylates an α-ketoacid and transfers the corresponding acyl-group onto a thiamine pyrophosphate (TPP) cofactor. This acyl-group is subsequently transferred to the E2 subunit (dihydrolipoyl acetyltransferase) which uses a prosthetic lipoic arm, covalently attached to a lysine residue, to transfer the acyl-group onto the thiol of coenzyme A (CoA), producing an acyl-CoA. As a result of this transfer, the lipoic arm becomes reduced to dihydrolipoamide. The E3 subunit (dihydrolipoamide dehydrogenase) uses its cysteine residues and FAD to transfer the electron from the dihydrolipoamide to NAD^+^, producing NADH and regenerating the disulfide bond in the lipoic arm for subsequent catalytic rounds.

The regulation of α-ketoacid dehydrogenases is important for numerous physiological processes including metabolic switching during fasting and feeding, response to hypoxia, and differentiation, and has broad relevance to many diseases including cardiovascular conditions, cancer, diabetes, and inborn error of metabolism.^1,5–14^ Such regulation is tightly controlled through layers of transcriptional, post-translational, and small metabolite-driven mechanisms. The most classic mechanisms include the inhibitory phosphorylations of the E1 subunit of PDHC and BCKDC, which is under the regulation by specific kinases and phosphatases,^15–18^ though similar phosphorylation is not known to regulate OGDC. Post-translational modifications of the E2 and E3 subunits, such as glutathionylation and cysteine nitrosylation, have also been reported to play significant regulatory roles.^19–22^

We recently found that in macrophages stimulated with lipopolysaccharide and interferon-γ (LPS+ IFNγ), PDHC and OGDC become profoundly inhibited.^23^ This inhibition drives substantial remodeling of the TCA cycle during the transition from the early pro-inflammatory stage to the later more suppressive stage, and has significant impacts on macrophage functions, in part, by dynamically controlling the level of itaconate and succinate^23^, two key metabolites with important immunoregulatory roles.^24–26^ We found that the inhibition of PDHC and OGDC is mainly driven by the significant decrease of the catalytically active lipoic arm on the E2 subunit.^16^ (Note: Here we use “lipoic arm” to refer to both the reduced and oxidized catalytically active lipoic moiety, and “lipoyl moiety” and “dihydrolipoyl moiety” are used to refer specifically to the oxidized and reduced forms of the lipoic arm, respectively.) However, the molecular mechanism controlling this decrease was unclear.

Here we show that PDHC and OGDC are regulated through novel covalent thiol modifications of the lipoic arm by reactive nitrogen species (RNS). Upon classical activation, macrophages produce a large amount of RNS by inducible nitric oxide synthase (iNOS) for pathogen killing.^27^ The RNS mediated modifications are efficiently delivered to the lipoic arm by CoA, the thiol-containing natural substrate for the E2 subunit. These modifications block the acyl-transferring activity of the lipoic arm, and cause profound inhibition of PDHC and OGDC, which can persist for an extended period. This work reveals a fundamental mechanism regulating lipoic arm-dependent enzymes, provides a new mechanistic link between RNS and mitochondrial metabolism, and has broad implications in many physiological and pathological conditions involving RNS production.

## Results

### Nitric oxide-dependent changes in PDHC and OGDC’s lipoic arms and activity

As shown in our previous work,^23^ LPS+ IFNγ stimulation of macrophages lead to a profound (>90%) reduction of PDHC and OGDC activity over time. This is in large part due to a decrease of the catalytically active lipoic arm, while total levels of PDHC and OGDC subunits do not decrease. We noticed that the decrease in the level of the active lipoic arm and enzyme activity temporally correlated with the increase in expression of iNOS in both the RAW 264.7 macrophage cell line (Fig 1a) and bone marrow derived macrophages (BMDM) (Fig 1b). Consistent with these changes, the intracellular levels of PDHC’s product acetyl-CoA and OGDC’s product succinyl-CoA also decreased over time (Fig 1c-d), in strong temporal correlation with the accumulation of intracellular citrulline, the product of iNOS (Fig 1e). Furthermore, when comparing across different stimulation conditions that induce varying levels of NO production in the two macrophage cell models, we found that higher iNOS activity (indicated by higher citrulline accumulation) correlated with less active lipoic arms on PDHC and OGDC (Supp figure 1). Together, these results demonstrate a quantitative correlation between nitric oxide (NO) production and the loss of PDHC and OGDC’s active lipoic arms and enzyme activity.

**Figure 1.**
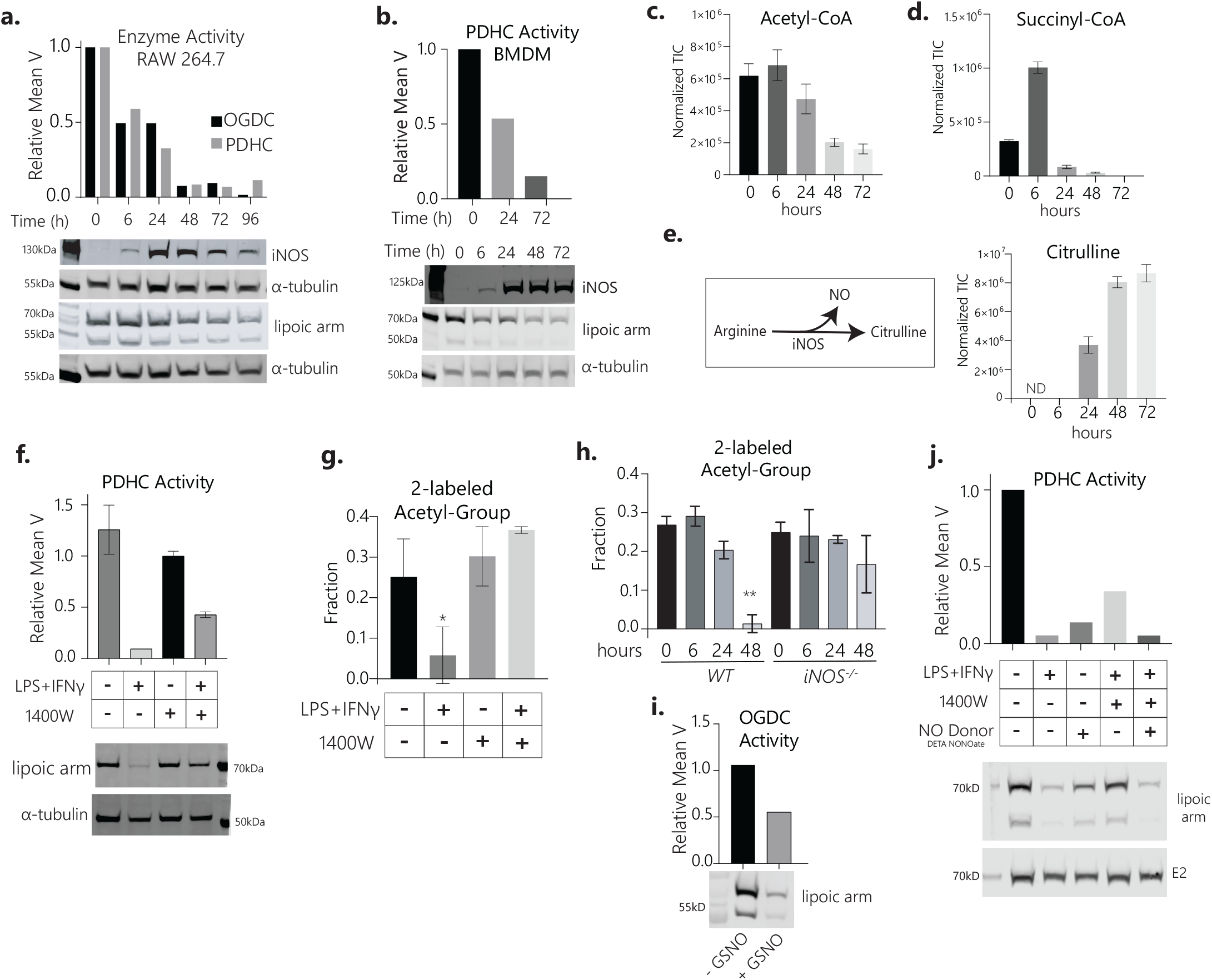
NO mediated inhibition of PDHC and OGDC via a mechanism involving lipoic arm. **a**. PDHC and OGDC activity, protein lipoylation state and iNOS expression in RAW 264.7 cells stimulated with LPS +IFN-γ for the indicated time. Experiments were performed independently twice with similar results. **b**. PDHC activity, protein lipoylation state and iNOS expression in BMDM stimulated with LPS +IFN-γ for the indicated time. Experiments were performed independently twice with similar results. **c-e**. The relative abundance of intracellular acetyl-CoA, succinyl-CoA and citrulline in BMDM stimulated with LPS +IFN-γ for the indicated time. Bar graphs with error bar represent mean ± s.d, n=3. **f**. The activity of PDHC (Mean ± SD, n=2) and lipoylation state of PDHC E2 subunit in the lysate of RAW 264.7 cells, unstimulated or stimulated with LPS+IFN-γ (48h), in presence or absence of iNOS inhibitor 1400W (40µM), as indicated. Western blot is representative of two independent experiments. **g**. The fraction of acetyl-CoA 2-labeled on the acetyl-moiety after incubation with U-^13^C-glucose tracer for 24h. Raw 264.7 cells were cultured ± LPS+IFN-γ stimulation ± iNOS inhibitor 1400W (40µM) for 48h. Bar graph represents mean ± sd, n=3. * indicates p<0.05 by t-test as compared to unstimulated untreated cells adjusted for multiple comparisons by Dunnett’s multiple comparison test. **h**. The fraction of acetyl-CoA 2-labeled on acetyl-moiety after incubation with 1,2-^13^C-glucose tracer in wild-type or iNOS knockout BMDMs stimulated with LPS+IFN-γ for indicated time. Bar graph represents mean ± sd, n=3. ** indicate p<0.0001 by t-test as compared to unstimulated WT cells (0h), adjusted for multiple comparisons by Dunnett’s multiple comparison test. **i**. The OGDC activity and lipoylation state in lysate of unstimulated RAW 264.7 cells with or without *in vitro* treatment of NO donor s-nitrosoglutathione (GSNO) (1mM for 3h). **j**. PDHC activity and protein lipoylation state in RAW 264.7 cells treated with indicated combination of stimuli (LPS+ IFNγ), iNOS inhibitor (1400W, 40µM), or NO donor (200µM DETA NONOate) for 48h.

To test the causal relationship behind this correlation, we inhibited iNOS in RAW 264.7 cells with the pharmacological inhibitor 1400W and found that it significantly prevented the decrease of PDHC’s active lipoic arm and enzyme activity (Fig 1f). To measure the intracellular flux through PDHC, we fed cells with ^13^C-glucose tracer and measured the fraction of acetyl-CoA labeled on the acetyl-moiety. We found that upon stimulation, acetyl-CoA labeling decreased substantially, indicating decreased intracellular PDHC flux, which is consistent with the decreased PDHC activity measured in cell lysate. The treatment with iNOS inhibitor fully rescued the intracellular flux through PDHC in stimulated macrophages (Fig 1g). Similarly, genetic knock out of iNOS also prevented the profound decrease in acetyl-CoA labeling from glucose in stimulated BMDM (Fig 1h), suggesting a rescue of intracellular flux. Inversely, treating cell lysate with an NO donor *in vitro* caused a decrease in the level of active lipoic arm and decreased OGDC activity (Fig 1i). In live cells, treating unstimulated macrophages with an NO donor caused a large decrease in active lipoic arm levels and PDHC activity, recapitulating what is observed in stimulated macrophages (Fig 1j). Additionally, the treatment of NO donor reversed the 1400W mediated protection of PDHC’s lipoic arm and activity in stimulated macrophages (Fig 1j). Together, these data show that the loss of catalytically active lipoic arm and the profound inhibition of PDHC and OGDC are largely mediated by NO.

### RNS cause thiol modifications on lipoic arm, inactivating its catalytic activity

We next investigated the molecular mechanism by which NO disrupts lipoic arm-dependent enzymes. The significant decrease in PDHC and OGDC activity was observed within a few hours of iNOS induction (Fig 1a), suggesting a fast-acting mechanism. We also observed that, when run on a gel, the E2 subunit shifted slightly towards higher molecular weight under conditions where NO level was high and the active lipoic arm level and PDHC activity were low (Fig 1j). Therefore, we hypothesized that RNS derived from NO can cause covalent modifications of the thiols on the lipoic arm, impairing its catalytic activity.

To evaluate the chemical reactivity of the lipoic arm to RNS, we incubated free lipoic acid (LA, C_8_H_14_O_2_S_2_) and its reduced form, dihydrolipoic acid (DHLA, C_8_H_16_O_2_S_2_), with NO donors, and found that DHLA reacted with RNS in solution and become depleted, while oxidized LA, which does not contain free thiol, did not react with RNS (Fig 2a-b). Untargeted LCMS analysis revealed that the reaction between DHLA and RNS yields a variety of products. The formulas of the most abundant products were identified based on their exact masses and isotopic distributions, and the likely structures were assigned based on literature knowledge of the chemical reactions between thiols and RNS.^28–30^ These products included: (1) at least two distinct species separated by their retention on chromatography with the formula C_8_H_15_O_3_S_2_N, which correspond to S-nitrosylation of either thiol of DHLA, a hydroxyl-disulfenamide, or a sulfinamide; (2) C_8_H_15_O_2_S_2_N, corresponding to a di-sulfenamide product; and (3) C_8_H_14_O_3_S_2_, corresponding to a thiosulfinate (Fig 2c).

**Figure 2.**
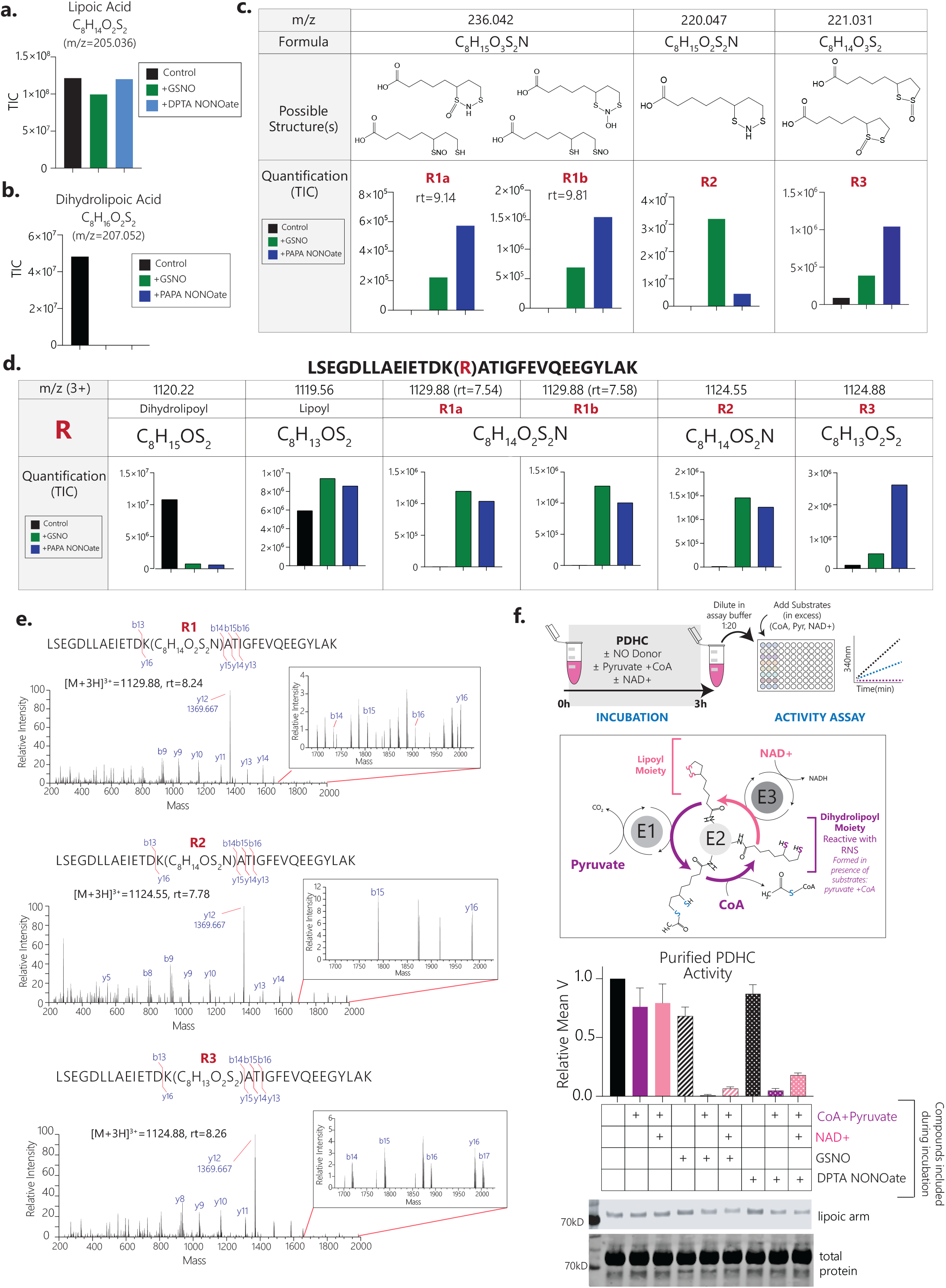
RNS cause thiol modifications on lipoic arm and inactivate its catalytic activity. **a**. Abundance of lipoic acid after incubation (3h, room temp) with or without indicated NO donor, GSNO or dipropylenetriamine NONOate (DPTA NONOate). **b**. Abundance of dihydrolipoic acid (DHLA) after incubation (3h, room temp) with or without indicated NO donor, GSNO or propylamine propylamine NONOate (PAPA NONOate). **c**. The appearance of major products, their formulas, possible structures, and abundances, after DHLA was incubated with indicated NO donor for 3h at room temperature. Unique identified products are designated by R# for clarity. M/z indicates exact mass or negative ion (single charge), rt indicates retention time. **d**. The depletion of substrate and appearance of indicated products after dihydrolipoyl peptide LSEFDLLAEIETDK(dihydrolipoyl)ATIGFEVQEEGYLAK was incubated with NO donors GSNO or PAPA NONOate in the presence of CoA for 3h at room temperature. M/z indicates exact mass of 3+ ion, rt indicates retention time, and R# corresponds to the identified lipoic modifications identified in figure 2c. **e**. MS/MS spectra of major products indicated in (d) confirming their identity. **f**. Activity and lipoylation state of purified PDHC measured following 3h incubation (RT) with NO donors (1mM GSNO or 1mM DPTA NONOate) in presence or absence of indicated substrates. After incubation with indicated compounds, enzyme activity was measured after dilution into assay buffer and addition of all substrates at a saturating level. Incubation of PDHC with CoA + pyruvate allow for formation of dihydrolipoyl moiety form of the lipoic moiety, which is reactive against RNS. Addition of NAD+ along with CoA + pyruvate during incubation allows PDHC’s lipoic arm to cycle between reduced and oxidized form through the full catalytical cycle. For the activity assay, bar graph with error bar represents mean ± s.d., n=4-5. **a-e**. Experiments were performed independently for at least two times with similar results.

We next examined whether similar reactions can occur non-enzymatically with a dihydrolipoylated peptide, by incubating NO donors with a synthesized E2 peptide containing a dihydrolipoyl group covalently attached to the lysine (K259, human; 258, mouse) residue [LSEGDLLAEIETDK(dihydrolipoyl)ATIGFEVQEEGYLAK]. We found that reaction with NO donors quickly depleted the dihydrolipoylated E2 peptide, and generated a series of thiol modifications, the same as those observed in the experiment with free DHLA, at significant amounts (Fig 2d,e). Incubation with NO donors also increased the amount of lipoylated peptide, suggesting RNS also promote the oxidation of dihydrolipoyl group to make a disulfide bond; however, the other thiol modifications collectively were more abundant (Fig 2d).

To test whether RNS mediated modifications of the lipoic arm can inhibit the activity of PDHC, we incubated purified PDHC with NO donors *in vitro*, and found that it nearly completely inactivated PDHC (Fig 2f). Importantly, such inactivation only occurred when PDHC’s substrates, pyruvate and CoA, were present with NO donors during the incubation period (Incubation with NO donors was performed for 3h prior to activity assay. During activity assay, substrates were added to all samples at saturating levels) (Fig 2f). The presence of pyruvate and CoA during incubation allows the conversion of PDHC’s lipoyl moiety to the more reactive dihydrolipoyl moiety through a partial catalytic cycle (Fig 2f). This dependence on pyruvate and CoA indicates that the RNS mediated inhibition specifically depends on modifications of the thiol of the lipoic arm, rather than another mechanism targeting different parts of the protein. PDHC was also profoundly inhibited by RNS when in the additional presence of the E3 subunit’s substrate NAD^+^, although the inhibition was slightly weaker than when PDHC was incubated with NO donors in the presence of just pyruvate and CoA (Fig 2f). This is expected, as the addition of NAD^+^ would allow PDHC to go through the complete catalytic cycle and effectively reduce the amount of time the lipoic arm stays in the reduced form with reactive thiols. Either in the presence of pyruvate + CoA or pyruvate + CoA + NAD, the inactivation of PDHC by NO donors was accompanied by decreased detection of active lipoic arm on E2 subunit (Fig 2f). Together, these experiments show that RNS can inhibit PDHC by causing covalent modification of the thiols on its lipoic arm.

### CoA delivers RNS-mediated modifications

Specificity and stoichiometry of post-translational modifications have long been key topics in protein research.^29,31,32^ As most known reactive oxygen and reactive nitrogen species-driven thiol modifications are non-enzymatic, it is important to understand how some modifications can be enriched sufficiently at functionally relevant sites to enable meaningful regulatory mechanisms. In this case, we found that the RNS-mediated inhibition of PDHC and OGDC can be nearly stoichiometric both *in vitro* and in cells. We hypothesized that the key factor behind such efficiency is CoA, the thiol-containing natural substrate for α-ketoacid dehydrogenases’ E2 subunits. It has been shown previously that low molecular weight (LMW) thiol-containing metabolites, particularly glutathione and CoA, play an important role in mediating cysteine nitrosylation in cells by forming LMW-SNOs. ^29,31,33,34^ Indeed, we found that, in presence of NO donors, CoA quickly became CoA-SNO (Supp Fig 2a). It is likely that, just like CoA, CoA-SNO can bind to the E2 subunit with high affinity and thus transfer the NO-modification onto the catalytic thiol of lipoic arm in a targeted manner, making this mechanism uniquely specific and efficient.

To test this hypothesis, we first examined the dependence of RNS-mediated PDHC inhibition on CoA. We found that while pyruvate alone would also allow generation of free thiols on the lipoic arm, incubating purified PDHC with an NO donor in the presence of pyruvate alone was not sufficient to cause inhibition; the presence of CoA was required (Fig 3a).

**Figure 3.**
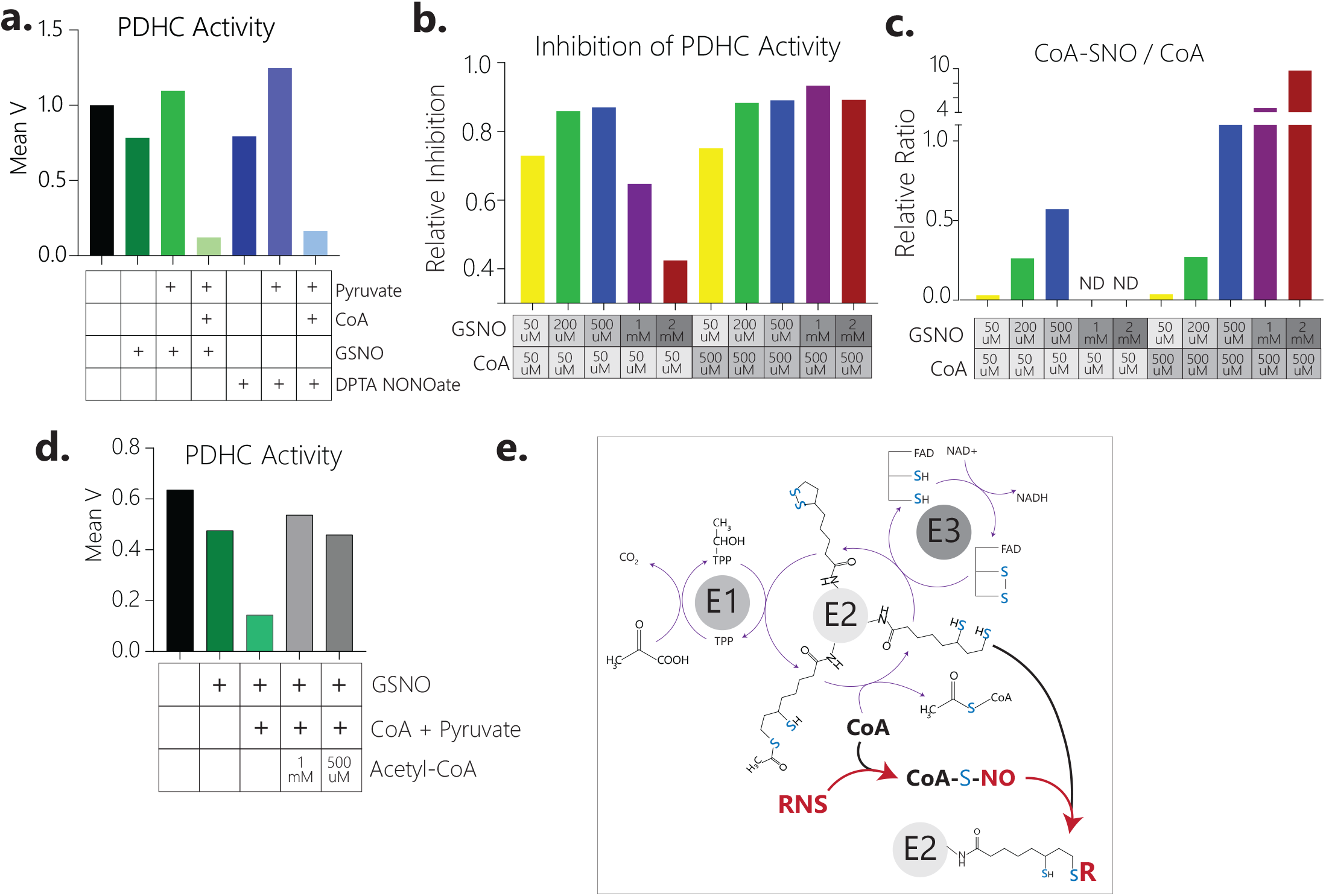
CoA delivers RNS modifications to the lipoic arm. **a**. Activity of purified PDHC after 3h incubation (at room temp) with NO donors GSNO (1mM) or DPTA NONOate (1mM), in presence or absence of CoA (5mM) and pyruvate (10mM). Trends have been repeated in another independent experiment with a different NO donor (PAPA NONOate) with similar results. **b**. Relative inhibition of PDHC when purified PDHC was incubated for 3h (RT) with indicated dosages of GSNO and CoA in presence of 500µM pyruvate. Relative inhibition is the loss of PDHC activity relative to PDHC control (incubated in buffer without substrate or NO donor for 3h). **c**. The relative ratio between CoA-SNO and CoA when indicated doses of GSNO and CoA are mixed in reaction as outlined in 3b. **d**. Activity of purified PDHC after 3h incubation ± GSNO ± substrates ± indicated amount of acetyl-CoA. Experiments were performed independently at least two times with similar results. **e**. PDHC enzymatic mechanism and model schematic. CoA, the thiol containing natural substrate of the E2 subunit, reacts with RNS and delivers the modifications to the lipoic arm, which inactivates its catalytical activity.

We next examined the efficiency of PDHC inhibition as a function of varying concentrations of NO donor and CoA. We found that the PDHC inhibition efficiency exhibited strong correlation with the ratio of CoA-SNO over CoA, as would be expected for a CoA-delivered modification mechanism, rather than the concentration of NO donor itself, as would be expected from a direct reaction between PDHC and RNS. Our LCMS analysis showed that, when CoA and the NO donor GSNO were mixed in solution, CoA-SNO is a major product and its formation increased with increasing concentration of NO donor, when the ratio between NO donor to CoA is relatively low; however, when the NO donor/ CoA ratio is very high, CoA-SNO formation decreases substantially (Supp Figure 2b), instead, the yield of other products (including CoA-disulfide, CoA-SG etc.) increases. We found that the dose-response curve of PDHC inhibition showed a similar biphasic trend: Under low CoA concentration (50 µM), increasing GSNO concentration from 50 to 500µM caused an increase in PDHC inhibition. However, further increasing GSNO concentration to 1mM and 2mM caused the inhibition of PDHC to become much weaker. In contrast, when CoA concentration was higher (500 µM), only the lowest dose of GSNO (50µM) caused a partial inhibition of PDHC, while concentrations of GSNO ranging from 200µM to 2mM each caused a near complete inactivation of PDHC (Fig 3b). Importantly, across all the examined conditions, the efficiency of PDHC inhibition strongly correlates with the ratio of CoA-SNO over CoA (Fig 3b,c), providing support for the CoA delivered modification mechanism.

Finally, if the binding of CoA-SNO to the E2 subunit is the key to targeting RNS-mediated modifications onto the lipoic arm, we would predict that the addition of acetyl-CoA, which can bind to the same site, would compete with CoA-SNO and protect PDHC activity. Indeed, this is what we observed. An excess amount of acetyl-CoA nearly completely prevented the loss of PDHC activity when incubated in the presence of NO donor and CoA (Fig 3d). These results strongly support that CoA delivers RNS mediated modifications onto the thiols of the E2 subunit’s lipoic arm (Fig 3e).

### The reversibility of RNS-mediated lipoic modifications

Here we found that RNS can cause a series of different thiol modifications on the lipoic arm (Fig 2c). Previous studies have shown that modifications such as S-nitrosylation are usually reversible by reaction with reducing agents such as glutathione and thioredoxin, while highly oxidized thiol modifications are generally irreversible.^35,36^ Some of the lipoic arm’s modification products have 5- or 6-member ring structures, which can also stabilize them further (Fig 2c). To test the chemical reversibility of the products of DHLA and RNS, we incubated the products with dithiothreitol (DTT), a common reducing agent. We found that the product C_8_H_15_O_3_S_2_N is mostly depleted upon incubation with DTT, while C_8_H_15_O_2_S_2_N and C_8_H_14_O_3_S_2_ are more resistant to the reduction by DTT (Supp Fig 3a), suggesting the reversibility varies based on the specific product.

To test the reversibility of enzyme modifications, we treated purified PDHC that had been incubated with NO donors, pyruvate, and CoA, with DTT. We found that DTT caused minimal recovery of the PDHC activity, suggesting that the inhibition is largely irreversible (Fig 4a). However, if NAD+ was included during the incubation period (in addition to NO donor, pyruvate and CoA), the resulting enzyme inhibition, although nearly as complete, was more easily reversed by DTT treatment (Fig 4a). This is likely because the presence of additional NAD^+^ causes a more reversible form of lipoic arm modification to become the dominant form, even though all the modification forms can effectively inactivate PDHC. Similarly, we treated cell lysates from unstimulated or stimulated macrophages, or lysates of unstimulated macrophages that had been incubated with NO donor *in vitro*, with DTT, and analyzed OGDC activity. We found that the loss of OGDC activity caused by both in cell stimulation and *in vitro* NO donor treatment was only slightly reversed by DTT treatment (Fig 4b).

**Figure 4.**
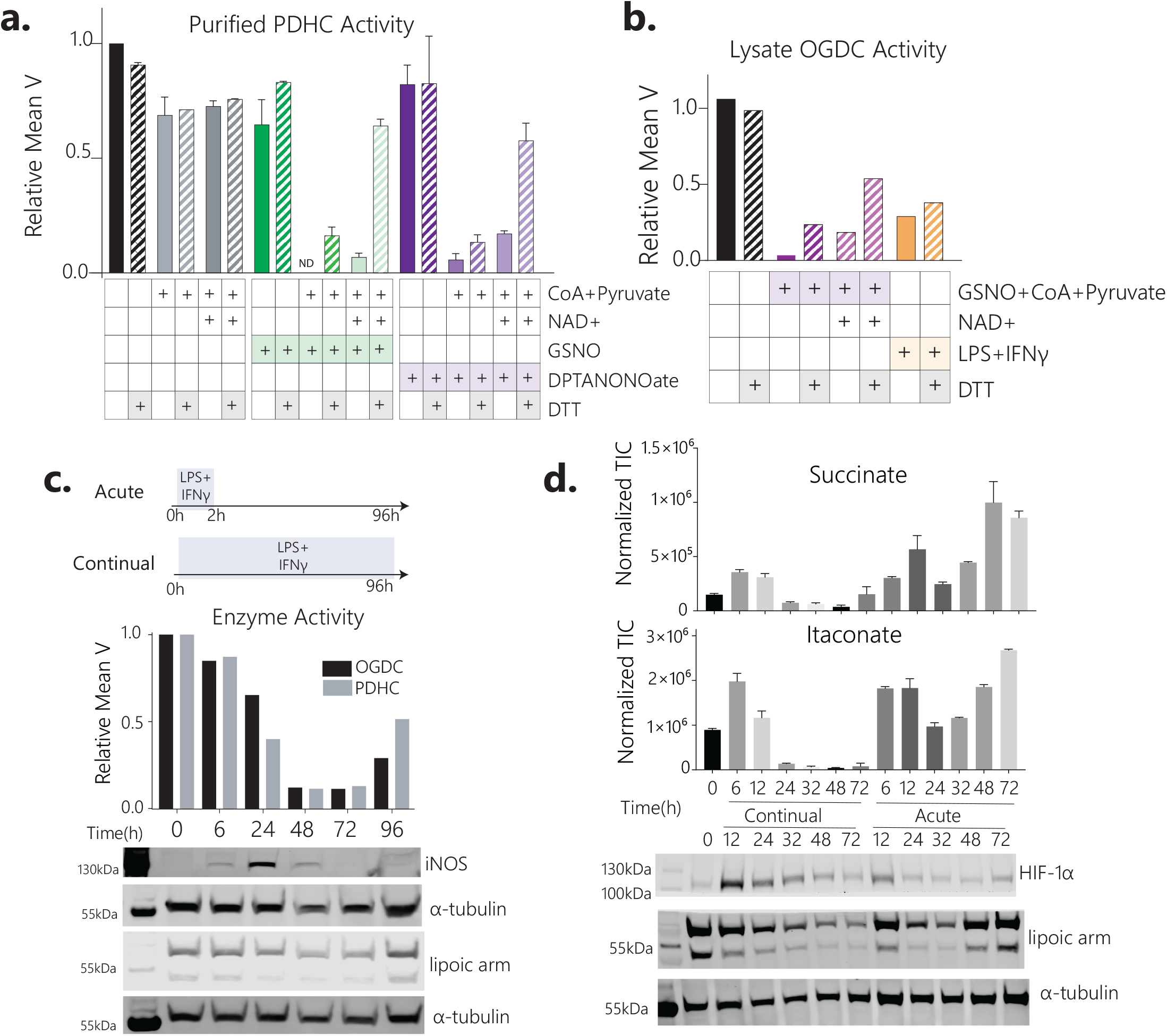
The reversibility of RNS-mediated PDHC and OGDC inhibition *in vitro* and in cells. **a**. Activity of purified PDHC after 3h incubation ± NO donors (GSNO or DPTA NONOate, 1mM) ± indicated substrates, followed by treatment of 20mM DTT or buffer control. Bar graph and error bars represent mean ± sd, n=2. **b**. Changes in OGDC activity in cell lysate upon *in vitro* DTT treatment (50mM, 30min). OGDC activity is measured in lysates of unstimulated RAW 264.7 cells which had been incubated *in vitro* with indicated combination of substrates (± 1mM CoA and 1mM pyruvate ± 1mM NAD+) and NO donor (± GSNO), before DTT treatment, and in lysate of LPS+IFNγ stimulated macrophages. Relative activity is normalized to OGDC activity in the lysate of unstimulated macrophages. **c**. PDHC and OGDC activity, lipoylation state, and iNOS expression in lysate of macrophages over a time course following 2h acute stimulation with LPS+IFNγ. **d**. The changes in intracellular succinate and itaconate level, lipoylation state, and HIF-1α level in RAW 264.7 cells over a time course either following 2h acute stimulation with LPS+IFNγ (Acute) or with continual exposure to LPS+IFNγ (Continual). For metabolite levels, bar graphs and error bars represent mean ± sd, n=3.

The reversibility and kinetics of PDHC and OGDC regulation can have a significant impact on cellular metabolism and function. In macrophages, upon acute stimulation with LPS+IFNγ for 2h, PDHC and OGDC activity and the unmodified (i.e. catalytically active) lipoic arm decreased substantially and remained very low through 48-72h post stimulation (Fig 4c). This long-lasting PDHC and OGDC inhibition, even after NO production had already diminished for over 24h, is consistent with the stability of the lipoic arm modifications we observed *in vitro*. However, by 96 hours post stimulation, the level of unmodified lipoic arm and PDHC and OGDC activity started to recover (Fig 4c). This contrasts with continual LPS+IFNγ exposure, where PDHC and OGDC inhibition is sustained throughout the time course, as continued NO production prevents the recovery of PDHC and OGDC (Fig 1a).

The dynamics of PDHC and OGDC regulation cause interesting downstream effects in macrophages upon activation. As we have shown previously,^23^ upon continual LPS+IFNγ stimulation, the TCA cycle undergoes a two-stage remodeling. The first stage is characterized by the accumulation of itaconate and succinate, which is driven in part by the fast upregulation of aconitate decarboxylase (IRG1). During the second stage, the inhibition of PDHC and OGDC shut off upstream flux into itaconate and succinate production, causing their levels to decrease (Supp Fig 3b). We now report that upon acute stimulation, while up to the first ∼32h the dynamic trends are similar to that observed upon continual stimulation, at very late time points, the eventual recovery of PDHC and OGDC’s lipoic arms and enzyme activity enables an intriguing second wave of itaconate and succinate accumulation (Fig 4d). Succinate and itaconate can stabilize HIF1α, a transcription factor with immunoregulatory functions, via competitive inhibition of prolyl hydroxylase. ^23,37–40^ Consistently, we observed that upon acute stimulation, HIF1α levels undergo two waves of accumulation, following the dynamic trends of succinate and itaconate abundance, and the lipoylation state of PDHC and OGDC (Fig 4d, Supp Figure 3c). This is in contrast to the single-wave of transient HIF1α accumulation seen upon continual stimulation, resulting from the sustained inhibition of PDHC and OGDC. These data demonstrate that RNS-mediated regulation of PDHC and OGDC through modifications of their lipoic arms can have important functional impacts in cells.

## Discussion

In this work we revealed RNS-mediated modifications of the lipoic arm as an important, previously unrecognized, mechanism to regulate mitochondrial a-ketoacid dehydrogenases PDHC and OGDC. We additionally illustrated that targeted delivery of RNS-mediated modifications by CoA makes this mechanism uniquely specific and efficient. This finding reveals a new biochemical link between RNS and cellular metabolism.

Several other mechanisms have been shown to regulate PDHC/OGDC, and to potentially link ROS and RNS with their regulation under different conditions and time scales.^21,41–44^ Interestingly, a recent report identified that the E3 subunit of PDHC can be nitrosylated, which inhibits its activity in activated macrophages, though the specific cysteine site was not identified.^20^ We propose that the E3 modification by NO is mechanistically connected to the lipoic arm modification revealed here. Because the normal function of the E3 subunit in the catalytic cycle is to re-oxidize the lipoic arm using a Cys-Cys active site (Fig 3e), it is likely that the E3 subunit can become modified on its catalytic cysteine via trans-nitrosylation from an NO-modified lipoic arm. To test this hypothesis, we analyzed the dependence of E3 modification on lipoic arm modifications in purified PDHC. We have established here that the modifications of lipoic arm by RNS requires the presence of substrates (pyruvate and CoA). We found that when PDHC is incubated with NO donors and substrates, especially when E3 subunit’s substrate NAD is also present, there is a small shift of E3 subunit towards higher molecular weight, indicating E3 modification. However, without substrates, NO donors do not cause such shift (Supp Fig 4). This result suggests that the modification of E3 subunit is promoted by RNS-mediated modifications of lipoic arm, rather than merely a direct reaction between E3 subunit and RNS. The connection between lipoic arm modification and E3 modification can further affect the efficiency, kinetics, and reversibility of RNS-mediated PDHC and OGDC inhibition.

Here we identified several types of covalent thiol-modifications that have not been reported on the lipoic arm before and showed that the reversibility of lipoic modifications can vary depending on the specific type of modification. The quantitative composition of different modifications under a specific condition depends on the ratio of RNS and CoA (and other thiols), the redox state, and potentially other chemical environmental factors such as pH. As such, variation in these conditions can impact the reversibility of RNS-mediated inhibition of PDHC and OGDC. The recovery of PDHC and OGDC we observed in macrophages after several days post-acute stimulation is likely a combined result of the reduction of reversible lipoic modifications, which would likely occur faster, and the general enzyme/lipoic arm turnover, which would take effect much more slowly. An important future direction is to quantitatively characterize the distribution of different forms of thiol modifications on lipoic arm, and their reactivity and recovery kinetics under various conditions. These studies would require developing a carefully validated LCMS-based proteomic method that sensitively and reliably quantifies the major lipoic modification forms, without perturbing the modifications during sample processing. Nevertheless, regardless of the specific distribution of different modifications, all identified RNS-driven modifications should block the lipoic arm from transferring the acyl-group and completing a normal catalytic cycle. Our combined data from various *in vitro* and in cell assays put forward strong evidence that these RNS-mediated modifications of the lipoic arm are an important mechanism for the regulation of PDHC and OGDC.

While this study mainly focused on PDHC and OGDC in macrophages, the implications can extend to a variety of cell types, including other cells that can produce NO, such as endothelial cells and neurons, as well as cells under the influence of NO production by neighboring cells.^29,31,45^ Besides PDHC and OGDC, BCKDC is another mitochondrial α-ketoacid dehydrogenase, which controls branched-chain amino acid oxidation. While BCKDC level in macrophages is low, it is highly expressed, and plays an important role, in other cell types such as adipocytes, myocytes and hepatocytes.^16,46–49^ The glycine cleavage system (GCS), which catalyzes a key step in mitochondrial one-carbon metabolism,^1^ also uses a similar lipoic arm-dependent catalytic mechanism. Regulation of these lipoic arm-dependent enzymes has been shown to be critical for energy homeostasis, proliferation, and various cellular functions, and have been implicated in numerous prevalent diseases, including diabetes, cancer, and heart failure.^49–53^ The discoveries presented here point to a new, broadly relevant mechanism for RNS to impact cellular physiology via the regulation of these key metabolic enzymes.

## Figure legends

**Supplementary Figure 1.**
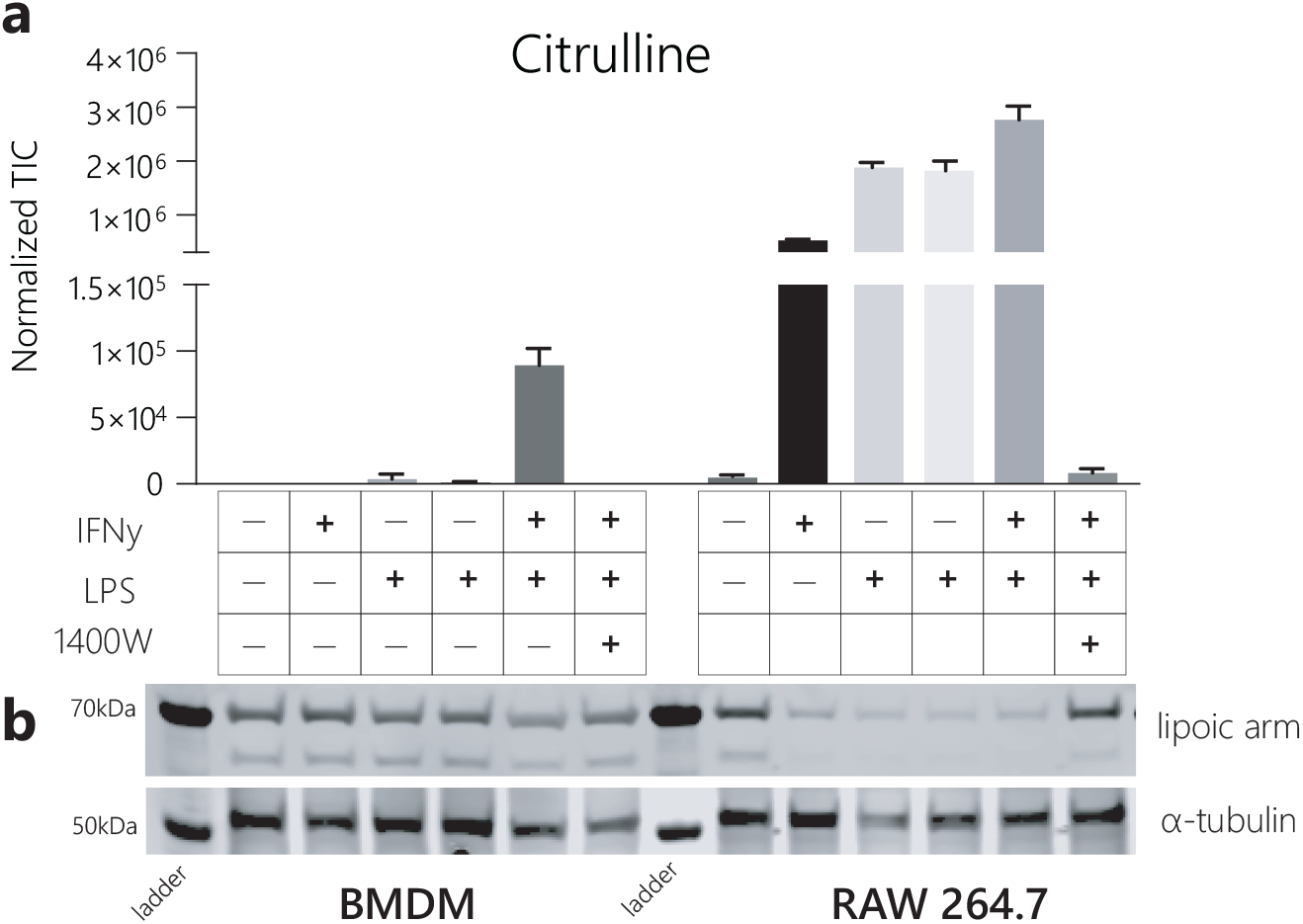
**a-b**. The correlation between intracellular accumulation of iNOS product citrulline (a) and the decrease of active lipoic arm (b) across different combination of stimuli (LPS and IFNγ) and iNOS inhibitor (1400W) between two different macrophage cell models, RAW 264.7 cells and murine bone marrow derived macrophages (BMDM). Bar graph and error bars represent mean ± sd, n=3.

**Supplementary Figure 2.**
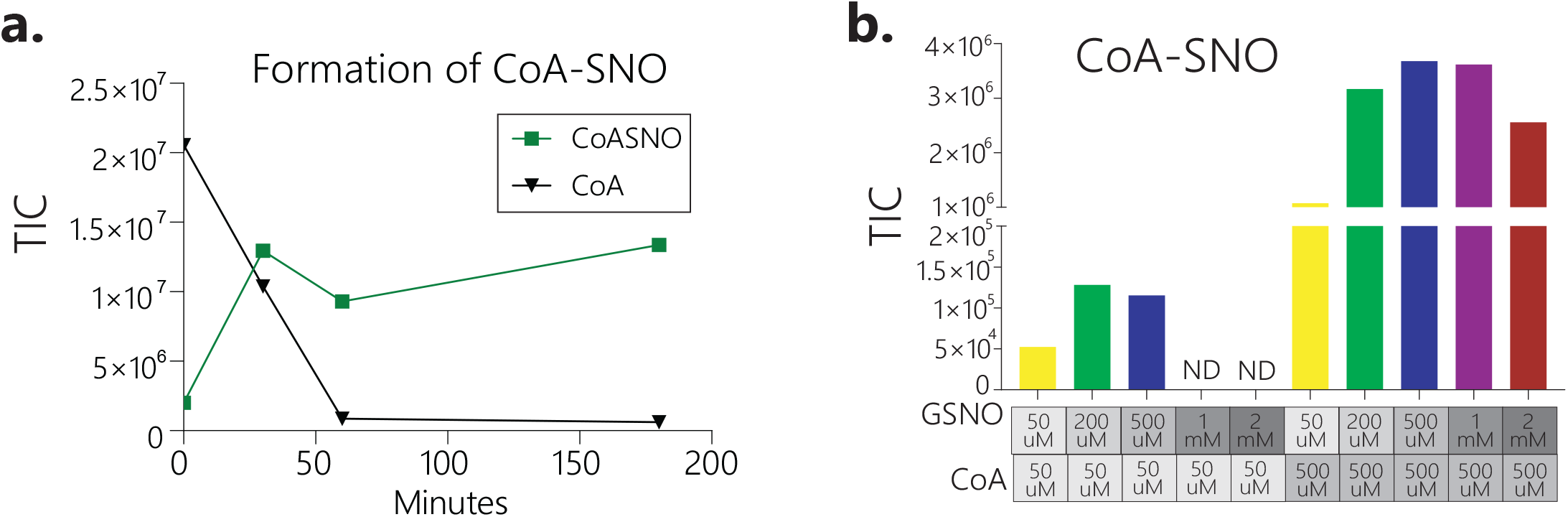
**a**. The decrease in CoA level and the appearance of CoA-SNO after CoA was mixed with NO donor PAPA NONOate in buffer for indicated time. **b**. CoA-SNO formed from non-enzymatic reaction between varying concentrations of CoA and GSNO, measured in the experiment shown in figure 3 b-c. ND indicates not detectable.

**Supplementary Figure 3.**
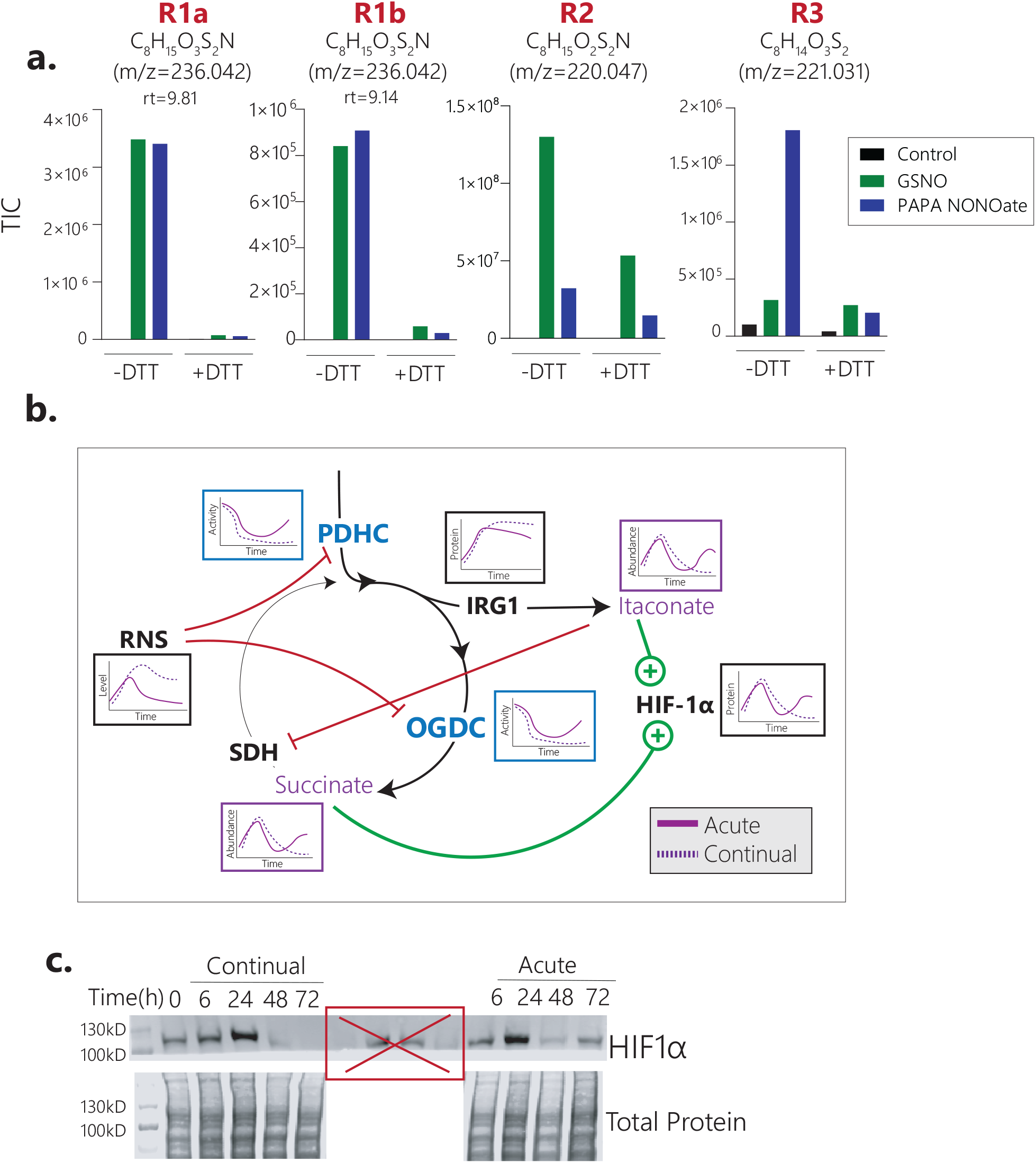
**a**. The major products formed as a result of incubation of dihydrolipoic acid (1mM, DHLA) with NO donors (1mM SNG or PAPA NONOate) and CoA (1mM), before and after incubation with DTT (10mM). Unique products are designated by R# for clarity and correspond to those shown in figure 2c. M/z indicates exact mass of negative ion (−1). **b**. Schematic showing the relationship between dynamic regulation of PDHC and OGDC and the changes in succinate, itaconate, and HIF-1α protein level in macrophages over a time course of acute or continual stimulation. Soon after stimulation, IRG1 expression increases, causing accumulation of its product itaconate. Itaconate can act as an inhibitor of succinate dehydrogenase (SDH), leading to a buildup of succinate. Both succinate and itaconate can inhibit prolyl hydroxylases and stabilize HIF1α, causing HIF1α to increase. However, as time progresses and RNS accumulate, PDHC and OGDC become inhibited, cutting off the influxes to itaconate and succinate, causing their decrease and thus destabilizing HIF-1α. At later time points upon acute stimulation, PDHC and OGDC activity recovers after iNOS expression subsides, while IRG1 level is still elevated, causing a second peak of itaconate, succinate and HIF-1α levels. This is in contrast with continual exposure to stimuli where iNOS expression remains high and PDHC and OGDC remain inhibited thus preventing the second peak of itaconate, succinate and HIF-1α levels. **c**. HIF-1α level in RAW 264.7 cells over a time course either following 2h acute stimulation with LPS+IFNγ (Acute) or with continual exposure to LPS+IFNγ (Continual). Crossed out part in the middle of gel are unrelated samples with different dosage of stimuli.

**Supplementary Figure 4.**
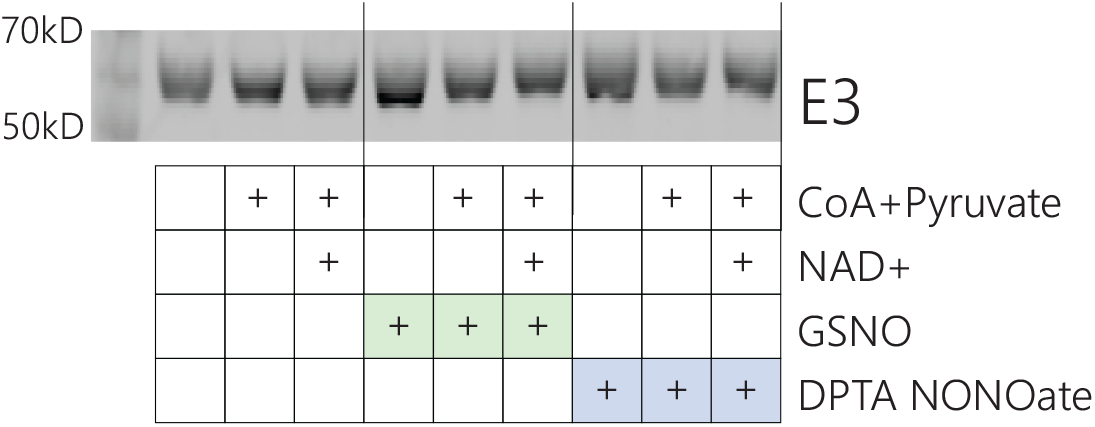
E3 subunit migration of purified PDHC after 3h incubation ± NO donors (GSNO or DPTA NONOate, 1mM) ± substrates (±1mM CoA and 1mM pyruvate, ± 6mM NAD+).

## Methods

### Cell Culture

RAW 264.7 cells (ATCC) were cultured in RPMI 1640 with 25mM Hepes, 1% penicillin/streptomycin and 10% FBS (dialyzed FBS was used for metabolite extraction experiments) at 37°C and 5% CO_2_. To stimulate the cells, 50ng/mL LPS (E. coli O111:B4, Sigma) and 10ng/mL IFN-γ (R&D Systems) were added to the media.

Murine bone marrow-derived macrophages (BMDMs) were isolated from wild-type C57Bl/6J mice or B6.129P2-Nos2tm1Lau/J (*iNOS*^*-/-*^) mice (Jackson Lab), as indicated. Mice were bred and maintained according to protocols approved by the University of Wisconsin-Madison Institutional Animal Care and Use Committee. Bone marrow cells were harvested from femurs and tibia of 6-12 week old mice (both male and females used). Cells were differentiated in RPMI 1640 containing 25mM Hepes, 1% penicillin/ streptomycin, 10% FBS and 20% L929 conditioned media. Three days post isolation, media were changed every day subsequently until 7 days post-isolation. Cells were then plated for experiments at 2.5*10^5^ cells/mL in RPMI 1640 containing 25mM Hepes, 1% penicillin/streptomycin, 10% FBS (dialyzed used for metabolite extraction experiments) and 20ng/mL M-CSF (R&D Systems). To stimulate BMDM, cells were incubated with 50ng/mL LPS (E. coli O111:B4, Sigma) and 30ng/mL IFN-γ (R&D Systems).

To avoid nutrient depletion, media for both RAW 264.7 cells and BMDMs were refreshed every day until collection. For continual stimulation experiments, LPS and IFN-γ were maintained in the media throughout. For acute stimulation experiments, cells were cultured with LPS and IFN-γ for 2h, after which stimuli were removed, and cells were washed with dPBS, and cell culture was continued in fresh media without stimuli for the remaining time-course.

In experiments where cells were treated with iNOS inhibitor or NO donor, the following chemicals were added: iNOS inhibitor, 1400W (Sigma, 40µM); NO donor Diethylenetriamine NONOate (DETA NONOate, Cayman Chemical, 200µM)

For experiments involving stable isotopic tracing, cells were incubated with media containing U-^13^C -glucose or 1,2-^13^C -glucose (Cambridge Isotopes), which replaces the unlabeled glucose in regular media at the same concentration, for 24h prior to collection. Data from labeling experiments was adjusted for natural abundance of ^13^C.

### Metabolite extraction

To extract intracellular metabolites, cells were washed three times with dPBS and metabolites were extracted with cold LCMS grade 80:20 methanol:H_2_O (v:v). Samples were dried under nitrogen flow and subsequently dissolved in LCMS grade water.

### LCMS analysis of small molecules

For the analysis of cellular metabolites or chemical reactions between small molecules (the reaction between lipoic acid or dihydrolipoic acid with RNS, or the reaction between CoA and RNS), samples were analyzed using a Thermo Q-Exactive mass spectrometer coupled to a Vanquish Horizon UHPLC. Analytes were separated on a 100 × 2.1 mm, 1.7µM Acquity UPLC BEH C18 Column (Waters), with a 0.2ml/min flow rate and with a gradient of solvent A (97:3 H_2_O:methanol, 10mM TBA, 9mM acetate, pH 8.2) and solvent B (100% methanol). The gradient is: 0 min, 5% B; 2.5 min, 5% B; 17 min, 95% B; 21 min, 95% B; 21.5 min, 5% B. Data were collected on a full scan negative mode. Setting for the ion source were; 10 aux gas flow rate, 35 sheath gas flow rate, 2 sweep gas flow rate, 3.2kV spray voltage, 320°C capillary temperature and 300 °C heater temperature.

Metabolites reported here were identified based on exact M/z and retention times determined with chemical standards. Data were analyzed with Maven.^63,64^ Relative metabolite levels were normalized to protein content.

### Calculation of acetyl moiety labeling in acetyl-CoA

The isotopic distribution of the acetyl moiety in acetyl-CoA, {M^Acetyl^}, was calculated based on measured isotopic distribution of acetyl-CoA{M^Acetyl-CoA^}, and coenzyme A, {M^CoA^} using the following equations:

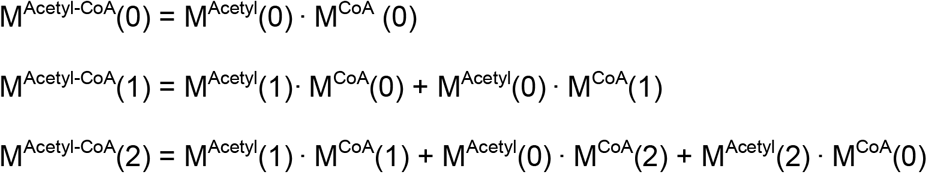

Where M^Acetyl-CoA^(i) and M^CoA^(i) are the measured fraction of acetyl-CoA and coenzyme A, respectively, with i labeled carbons, after incubation with ^13^C glucose tracer. Labeling of acetyl moiety in acetyl-CoA was calculated by fitting M^Acetyl^(0), M^Acetyl^(1), and M^Acetyl^(2) based on these equations. In cases where CoA signal was not good enough, ATP labeling patterns were used to impute CoA labeling data prior to acetyl-CoA fitting.

### Activity assays of PDHC and OGDC in cell lysate

Pyruvate dehydrogenase complex activity in cell lysate was analyzed using a PDH enzyme activity microplate assay kit (Abcam), as per manufacturer’s instructions. This kit measures PDHC activity by monitoring pyruvate-dependent NADH production. The NADH level was measured by absorbance of NADH coupled dye (450nm) using a BioTek^®^ Epoch2 plate reader.

To evaluate OGDC activity, mitochondria were isolated and the α-ketoglutarate-dependent NADH production rate in mitochondria lysate was measured by monitoring NADH absorbance at 340nm, according to a previously established method.^23^ Cells were lysed in hypotonic lysis buffer (20mM Tris buffer pH=7.4, 1mM EDTA with phosphatase and protease inhibitors (Pierce™ Protease and Phosphatase Inhibitor Tablets)) and incubated on ice for 30 min. Lysate was then homogenized in a dounce homogenizer and then separated via centrifugation at 700 × g, 4°C, for 10 min, in a mannitol sucrose solution (440mM mannitol/140mMsucrose, 20mM Tris, 1mM EDTA, pH 7.4). The mitochondrial fraction was resuspended in RIPA buffer with protease and phosphatase inhibitors (ThermoFisher). Mitochondrial lysate (∼30µg protein) was diluted in 200µL of assay buffer (1mM MgCl, 400µM thiamine diphosphate, 50mM HEPES, 10% glycerol, 0.05% BSA) and combined with substrates (α-ketoglutarate, CoA, NAD); absorbance at 340nM was measured continously using a BioTek® Epoch2 microplate reader.

To investigate the ability of nitric oxide to inhibit PDHC/OGDC activity in isolated lysate, cell lysate (isolated using methods above) was incubated with nitric oxide donors at room temperature prior to activity measurement. The following NO donors were used: S-nitrosoglutathione (GSNO, Cayman Chemical) and Dipropylenetriamine NONOate (DPTA NONOate, Cayman Chemical). The reversibility of the RNS-mediated PDHC inhibition was tested by treating the lysate with excess amount of DTT (50mM) for 30min at room temperature after incubation with NO donors was complete.

### In vitro assays of purified PDHC

Purified porcine PDHC (Sigma) was used for *in vitro* assays of RNS-mediated inhibition of PDHC. Enzyme (20-100mU/mL) was incubated at room temperature with NO donors, GSNO, DPTA NONOate and PAPA NONOate (Cayman Chemicals) along with combinations of CoA, NAD+, pyruvate, and acetyl-CoA, as indicated in figure legends, in 20mM NaPO_4_ buffer (pH=7.2) for 3h. In experiments testing the reversibility of the RNS-mediated PDHC inhibition, reactions were subsequently incubated with excess amount of DTT (20mM) or buffer control for 30min at room temperature.

At the end of incubation, the reaction solution was further diluted 1:20 in assay buffer (20mM NaPO_4_ buffer) and combined with an assay solution containing 100uM TPP, CoA (1-5mM), pyruvate (1-5mM) and NAD+ (2.5-50mM). The concentrations of the substrates in assay conditions were chosen so that (1) all substrates are at saturating levels so the maximal enzyme activity is measured; (2) the substrates added in assay buffers are in excess of any carried over substrates added during incubation period (after the 1:20 dilution), so the variation of final substrate concentrations in assay condition across samples are negligible (3) CoA level added in assay condition is in excess of any leftover NO donor, so possible consumption of CoA by reaction with RNS would not significantly affect the assay. This was further verified by LCMS analysis of the reaction mix.

PDHC activity was measured by the rate of NADH absorbance (340nM) increase using a BioTek® Epoch2 microplate reader. NADH production was read continuously, and mean V values were calculated from a linear portion of the curve (up to first 10-30min depending on the experiment) and normalized to protein content. In those experiments where large amount of acetyl-CoA was incubated with purified enzyme to test its protection against CoA-SNO mediated inhibition, varying levels of acetyl-CoA were added back to the assay solutions after the incubation period, such that in the assay condition, the concentration of acetyl-CoA (a potential inhibitor of PDHC) were even across samples.

### Analysis of the reaction between LA / DHLA and RNS

Lipoic acid (Cayman Chemical) or dihydrolipoic acid (DHLA, Sigma-Aldrich) was incubated with the NO donors, GSNO, DPTA NONOate, and PAPA NONOate (Cayman Chemical) at room temperature in 20mM NaPO_4_ reaction buffer for 1h or 3h. For reversibility test, reaction mixes were then subsequently incubated with 10mM DTT or buffer control at room temperature. At the end of incubation, reactions samples were diluted in LCMS grade methanol and immediately injected onto the LCMS using the same LCMS method outlined in ***LCMS analysis of small molecules***.

To identify major products from the reaction between DHLA and RNS, LCMS features were pulled from untargeted analysis of control (DHLA incubated without NO donor), and NO donor-treated samples. The features were compared to find major peaks present in NO donor-treated samples but not in control, and these peaks were clustered based on retention time and chromatographic profile to identify groups of unique chemical species and their parent peak (because each compound can give many LCMS features that are isotopic peaks or adducts of the parent compound). The formulas were identified based on exact mass and the isotopic distribution, especially the natural S34 peak and C13 peaks that indicate their composition.

### Analysis of reaction between lipoylated E2 peptide and NO donor

Dihydrolipoylated E2 peptide, LSEGDLLAEIETDK(dihyrdrolipoyl)ATIGFEVQEEGYLAK (>90% purity) was synthesized by GenScript. Because some of the dihyrdrolipoyl moiety would spontaneously oxidize to lipoic moiety in solution, to reduce the disulfide bond, prior to incubation with NO donors, the peptide (50uM in 20mM NaPO4 buffer) was reduced using 25mM TCEP (70C for 20min), and then TCEP was removed by Stage-Tip. Briefly, stage-tips were assembled by layering C18 Empore reversed phase extraction disks, 3M into the bottom of 200uL tip using a 16-guage needle and plunger. Tips were conditioned with 100uL of 100% ACN (2000xg 1min) followed by 100µL 0.5% TFA (2000xg 1min). Reduced peptide sample was added to top of the stage tip and then centrifuged at 500xg for 2min. The stage-tip was then washed 4x with 150µL of 0.5% TFA/5%ACN (2000xg 2min), and the sample was then eluted using 100µL of 70%ACN/0.5% TFA (2000xg 3min). Sample was then diluted in reaction buffer (20mM NaPO_4_ buffer containing 3mM CoA). NO donors (GSNO or PAPA-NONOate) were added to the solution and incubated at room temperature for 1.5h, after which samples were analyzed by LCMS.

Peptides were separated on a 100 × 2.1 mm, Accucore Vanquish UHPLC C18+ column (Thermo), with a 0.25ml/min flow rate and with a gradient of solvent A (5:95 H_2_O:ACN) and solvent B (0.1% formic acid). The gradient is: 0 min, 90% B; 2 min, 90% B; 14 min, 1% B; 15 min, 1% B; 16 min, 90% B; 20 min, 90% B. Data were collected on a positive mode with full scan/ddMS2 (loop count: 3, isolation window: 2 m/z, fragmentation with a stepped NCE: 20, 25, 30). Data were analyzed by Xcalibur.

### Gel electrophoresis and immunoblotting

Whole cell lysate samples were prepared by lysing cells in RIPA buffer with phosphatase and protease inhibitors (Pierce™ Protease and Phosphatase Inhibitor Tablets) after washing with dPBS three times. Protein content of lysate was quantified using BCA assay kit (ThermoFisher). Whole cell lysates or purified PDHC reactions were separated on 4-12% Bolt Gel (ThermoFisher) after heat denaturation in sample buffer (ThermoFisher). No reducing buffer was added to samples to minimize potential perturbations of thiol modification on proteins. Proteins were subsequently transferred to 0.2µM nitrocellulose membrane. Membranes were blocked in 5% nonfat milk in TBS-T buffer and probed with the following primary antibodies: anti-HIF-1α (CST 14179), anti-α-tubulin (Abcam ab7291), anti-PDH E2 (Abcam ab66511), anti-lipoic acid (Calbiochem, 467695), anti-iNOS (ThermoFisher PA5-17100), anti-PDH E3 (Abcam ab133551). Primary antibody incubation was followed by incubation with Li-Cor® secondary antibodies (goat anti-rabbit 800CW, goat anti-mouse 680RD). Total protein was quantified prior to blocking using Revert Total Protein Stain (Licor). Membranes were imaged with a Li-Cor® Odyssey ClX and quantified using ImageStudio™ Lite software.

## Acknowledgements

We thank Dr. Michael Marletta at University of California, Berkeley for helpful discussions. Gretchen Seim is supported by NRSA Individual Predoctoral Fellowship F31AI152280.

## Competing interest statement

The authors declare no competing interests.

## Notes

### Competing Interest Statement

The authors have declared no competing interest.

